# Age-Related Type I Collagen Modifications Reveal Tissue-Defining Differences Between Ligament and Tendon

**DOI:** 10.1101/2021.03.25.437061

**Authors:** David M. Hudson, Marilyn Archer, Jyoti Rai, MaryAnn Weis, Russell J. Fernandes, David R. Eyre

## Abstract

Tendons and ligaments tend to be pooled into a single category as dense elastic bands of collagenous connective tissue. They do have many similar properties, for example both tissues are flexible cords of fibrous tissue that join bone to either muscle or bone. Tendons and ligaments are both prone to degenerate and rupture with only limited capacity to heal, however; healing outcomes for tendons are often faster than ligament injuries. Type I collagen constitutes about 80% of the dry weight of tendons and ligaments and is principally responsible for the core strength of each tissue. Collagen synthesis is a complex process with multiple steps and numerous post-translational modifications including proline and lysine hydroxylation, hydroxylysine glycosylation and covalent cross-linking. The chemistry, placement and quantity of intramolecular and intermolecular cross-links are believed to be key contributors to the tissue-specific variations in material strength and biological properties of collagens. As tendons and ligaments grow and develop, the collagen cross-links are known to chemically mature, strengthen and change in profile. Accordingly, changes in cross-linking and other post-translational modifications are likely associated with tissue development and degeneration. Using mass spectrometry, we have compared tendon and ligaments from fetal and adult bovine knee joints to investigate changes in collagen post-translational properties. Although hydroxylation levels at the type I collagen helical cross-linking lysine residues were similar in all adult tissues, ligaments had significantly higher levels of glycosylation at these sites compared to tendon. Differences in lysine hydroxylation were also found between the tissues at the telopeptide cross-linking sites. Total collagen cross-linking analysis, including mature trivalent cross-links and immature divalent cross-links, revealed unique cross-linking profiles between tendon and ligament tissues. Tendons were found to have a significantly higher frequency of smaller diameter collagen fibrils compared with ligament, which we propose is a consequence of the unique cross-linking profile of each tissue. Understanding the specific molecular characteristics that define and distinguish these specialized tissues will be important for designing improved orthopedic treatment approaches.

## Introduction

Tendons and ligaments are collagenous connective tissues that are often grouped as the tough, flexible cords that hold together the musculoskeletal system. Indeed, the only consistent distinction made between the two tissues in much of the literature is that tendons join muscle-to-bone whereas ligaments join bone-to-bone. Tendons and ligaments do have many general similarities including biomechanics (high tensile strength) and composition (predominantly type I collagen) and the common public health issue that half of all musculoskeletal injuries are the result of ligament or tendon damage [1]. Despite the prevalence of tendon and ligament injuries and the economic cost of surgical repair, there remains a fundamental lack of understanding of tendon and ligament biology as it relates to development and healing [2]. For example, it is not fully known why injured intra-articular ligaments, such as anterior cruciate ligament (ACL) and posterior cruciate ligament (PCL), have no significant healing capacity and often require surgery [3], whereas vascularized tendons and ligaments, such as medial collateral ligament (MCL) and quadriceps tendon (QT), have a limited ability to heal [2,4]. Although some tendon and ligament injuries such as strains and sprains may not require surgery, the healing process involves the formation of scar tissue that may require days, months and even years to remodel [2]. Furthermore, remodeled MCL has been shown to be weaker, less stiff and absorb less energy compared to normal MCL [5]. Even after surgical repair, tendons often fail to regain their native structure and function, which can lead to re-tear events [6]. Unlike surgical repair, which is defined as any surgical procedure attempting to restore the original injured tissue; surgical reconstruction is a surgical procedure that replaces the injured tissue with a graft [3]. For example, in the case of ACL reconstruction, the autografted tissue is often obtained from a tendon source. However, it is unclear if mature tendon is the best substitute for ligament reconstruction. Recognizing tendons and ligaments as distinct specialized tissues with distinguishing features could help inform future applications and improvements in orthopedic outcomes.

Interestingly, it has been shown that injured fetal tendons have the capacity to regenerate fully thus restoring native structure and function [7]. This has been correlated to embryonic tendons having a higher cell density and activity, both of which decrease as tendons develop and mature [8]. Yet the specific conditions, which enable fetal regeneration and scarless healing, have not been fully explored. Furthermore, it is unknown if this capacity for healing in fetal tendons can be applied to adult tendons. Defining any collagen post-translational variances, such as cross-linking, between fetal and adult tendons and ligaments may potential help provide insight into differences in tissue remodeling and healing. For example, during growth and development, both tendons and ligaments have immature and reversible collagen cross-links that, with age, increase in strength and form insoluble and permanent cross-links [5]. Collagen cross-linking is the definitive post-translational modification that imparts tensile strength to connective tissues.

Both tendons and ligaments are predominantly composed of type I collagen (approximately 80% by dry weight). Type I collagen biosynthesis includes many post-translational modifications such as hydroxylation (e.g. 4-hydroxyproline, 3-hydroxyproline, 5-hydroxylysine), glycosylation (e.g. galactosyl-hydroxylysine, glucosylgalactosyl-hydroxylysine) and cross-linking (e.g. hydroxylysine pyridinoline) [9]. The importance of collagen post-translational modifications is underscored by the many heritable disorders that can result in altered collagen biosynthesis [10,11]. A careful understanding of the tissue-specific nature of collagen post-translational modifications is essential for distinguishing between type I collagens across connective tissues. For example, tendon type I collagen is known to contain more 3-hydroxyproline modifications than type I collagen from skin or bone [12,13]. Whereas, hydroxylysine glycoside content in tendon type I collagen is significantly less than that of skin or bone [14]. In fact, tendon type I collagen is particularly limited in glycosylations, with only one site, a hydroxylysine at K87, in each α-chain that appears to be consistently glycosylated. Additional partial glycosylation sites in tendon and ligament include K174 and K219 in both α-chains, but only at low occupancy [15,16]. The K87 residue is of unique importance as it participates in cross-link formation with α1(I) C-telopeptide aldehydes (likewise helical α1(I)K930 and α2(I)K933 are the substrates for N-telopeptide aldehyde cross-link formation) [17]. Although, the functional effect of glycosylation in collagens is not fully understood, a broad spectrum of roles have been proposed including facilitating matrix remodeling [18,19], collagen fibrillogenesis [20–22], bone mineralization [23], and participating in collagen cross-linking [24–27].

In the current study, we set out to define distinct tissue specific features that may distinguish tendon and ligament in development. Recent studies focusing on collagen post-translational modifications in ligaments have been carried out on periodontal ligament [28–30]. In this study, we characterize the collagen profile of multiple ligament tissues isolated from adult and fetal bovine knee joints. Our focus was to compare type I collagen post-translational variances between intra-articular ligaments (PCL and ACL), vascularized ligaments (lateral collateral ligament (LCL) and MCL) and tendon (QT). We propose that modifications unique to ligament type I collagen, including collagen cross-links, have evolved to enable the matrix to adapt and function in a non-vascularized environment.

## Results

### Tendon and ligament collagen have similar prolyl 3-hydroxylation fingerprints

Tendon (QT) and ligaments (ACL, PCL, LCL and MCL) were harvested from hindlimb knee joint of 18-month (adult; n=3) and fetal bovine animals (n=3). Collagens were extracted from adult and fetal bovine tendons and ligaments by acid extraction. Although all tissues produced a similar banding pattern of collagen α-chains on SDS-PAGE, the adult tissue collagens were consistently less extractable compared to fetal tissue (Figure 1). Type I collagen was predominantly extracted from all tissues. Mass spectrometric analysis of the collagen α-chains revealed distinct developmental post-translational features between adult and fetal tendons and ligaments. We initially focused on the known prolyl 3-droxylase 2 (P3H2)-catalyzed 3-hydroxyproline modifications, which in type I collagen are the α1(I)P707 site and the C-terminally located (GPP)n motifs. Type I collagen isolated from adult tendon is known to contain higher levels of prolyl 3-hydroxylation compared to other connective tissues, such as bone and skin [13,31]. This molecular phenotype, which was once thought unique to tendon, was also observed across all investigated adult bovine ligaments (Figure 2, Table 1). Another commonality found between the two connective tissues was the post-translational increase in prolyl 3-hydroxylation that occurs during development. Fetal tendons have previously been shown to exhibit only minimal levels of the prolyl 3-hydroxylation in type I collagen compared to adult tendons [31–34]. It was consistently observed in the current study that the P3H2-catalyzed sites were minimally occupied in fetal tendons and ligaments (Table 1). Figure 2 highlights the difference in prolyl 3-hydroxylation in the (GPP)_n_ motif between adult and fetal ACL as seen by MS analysis. The known prolyl 3-hydroxylase 1 (P3H1)-catalyzed 3-hydroxyproline sites (α1 (I)P986 and α2(I)P707) were almost fully occupied in both adult and fetal ligaments and tendons (Table 1).

**Figure 1.**
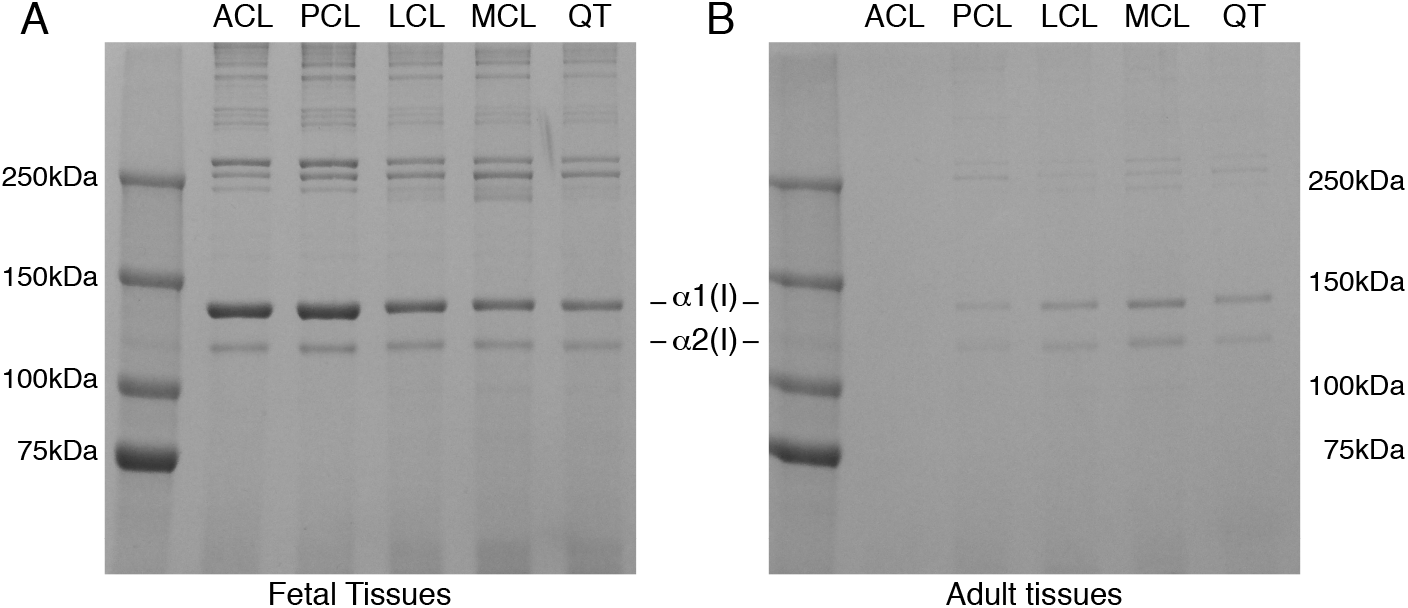
SDS-PAGE reveals a decrease in collagen extractability in adult tendons and ligaments compared to fetal tissue. Acid labile aldimine cross-links are broken with mild acetic acid treatment, allowing native type I collagen monomers to be extracted from the tissue. Collagen was more acid extractable from fetal tissues (A) than adult tissues (B). Anterior cruciate ligament (ACL), posterior cruciate ligament (PCL), lateral collateral ligament (LCL), medial collateral ligament (MCL), Quadriceps tendon (QT). The reduction in collagen extractability from adult tissues is consistent with an increase in mature collagen cross-links with development.

**Table 1.** Comparison of 3Hyp occupancy in type I collagens from adult and fetal bovine tendon and ligaments. Mean percentage of 3Hyp at each major substrate site in type I collagen from tendons and ligament tissues (n=3). The percentages were determined based on the ratios between the ion-current yields of each post-translational variant as previously described.

| | $\alpha 1(I)P986$ | | | | $\alpha 1(I)P707$ | | | | $\alpha 2(I)P707$ | | | | $\alpha 1(I)GPP$ | | | |
| --- | --- | --- | --- | --- | --- | --- | --- | --- | --- | --- | --- | --- | --- | --- | --- | --- |
|  | Q | A | L | M | Q | A | L | M | Q | A | L | M | Q | A | L | M |
| Adult<br>(n=3) | 95% | 96% | 95% | 95% | 60% | 43% | 52% | 62% | 92% | 90% | 99% | 96% | 57% | 50% | 53% | 62% |
| Fetal<br>(n=3) | 96% | 96% | 96% | 96% | 19% | 21% | 21% | 25% | 91% | 98% | 97% | 98% | 8% | 7% | 7% | 8% |

### Type I collagen cross-linking lysines (K87) from ligaments are highly glycosylated

Using in-gel trypsin digestion we investigated the post-translational quality of the type I collagen helical cross-linking residues (α1(I)K87 and α2(I)K87) and the C-telopeptide cross-linking residue (α1 (I)K1030) in each connective tissue. The cross-linking K87 residues from both α-chains of type I collagen from fetal ligament and tendon were fully hydroxylated and subsequently glycosylated (glucosylgalactosyl-hydroxylysine) (Figure 3). A distinct feature of tendon (QT) development compared to ligament (ACL) was the significant decrease in glycosylation at the K87 residues in adult tissue (Figures 3 and 4). The adult ligaments investigated in this study varied significantly in the degree of modification at the K87 residues. It appears that the intra-articular ligaments (ACL) retain a higher level of glycosylation in adulthood compared to vascularized connective tissues (QT, LCL, MCL). These results are summarized in Table 2. Collagen from the intra-articular ligaments, ACL (Figure 3) and PCL (not shown), had comparable post-translational modification profiles at α1(I)K87 and α2(I)K87. The post-translational quality of α1(I)K930 and α2(I)K933 could only be detected using bacterial collagenase digestion followed by resolution of peptide fragments by reverse phase chromatography. Although this approach is not as quantitative as in-gel-trypsin, our data support almost 100% hydroxylation of both α1(I)K930 and α2(I)K933 in fetal and adult tendons and ligaments (Table 2).

**Figure 2.**
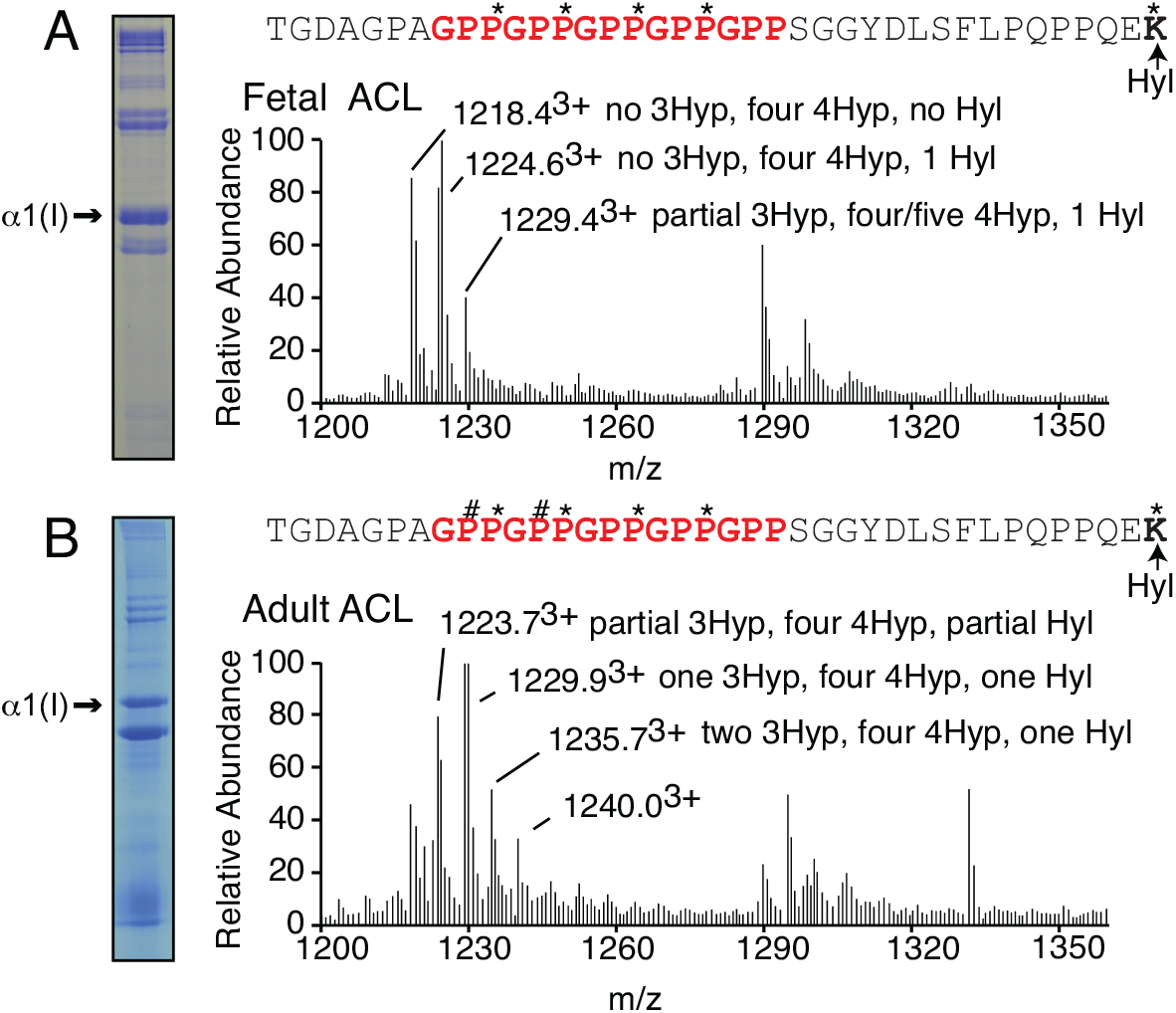
Type I collagen from ligaments exhibits age-dependent 3-hydroxyproline profile. LC-MS profiles of in-gel trypsin digests of the collagen α1(I) chain from fetal and adult bovine ACL. (A) MS profile of fetal bovine ACL shows only partial 3-hydroxylation (~8%) at the α1(I) C-terminal GPP motif; (B) MS profile at the same site of adult bovine ACL shows a hydroxylation like bovine tendon 3-hydroxylation (~60%). The trypsin digested peptide is shown with P# indicating 3Hyp, P* indicating 4Hyp and K* indicating Hyl. See Table 1 for more details.

**Figure 3.**
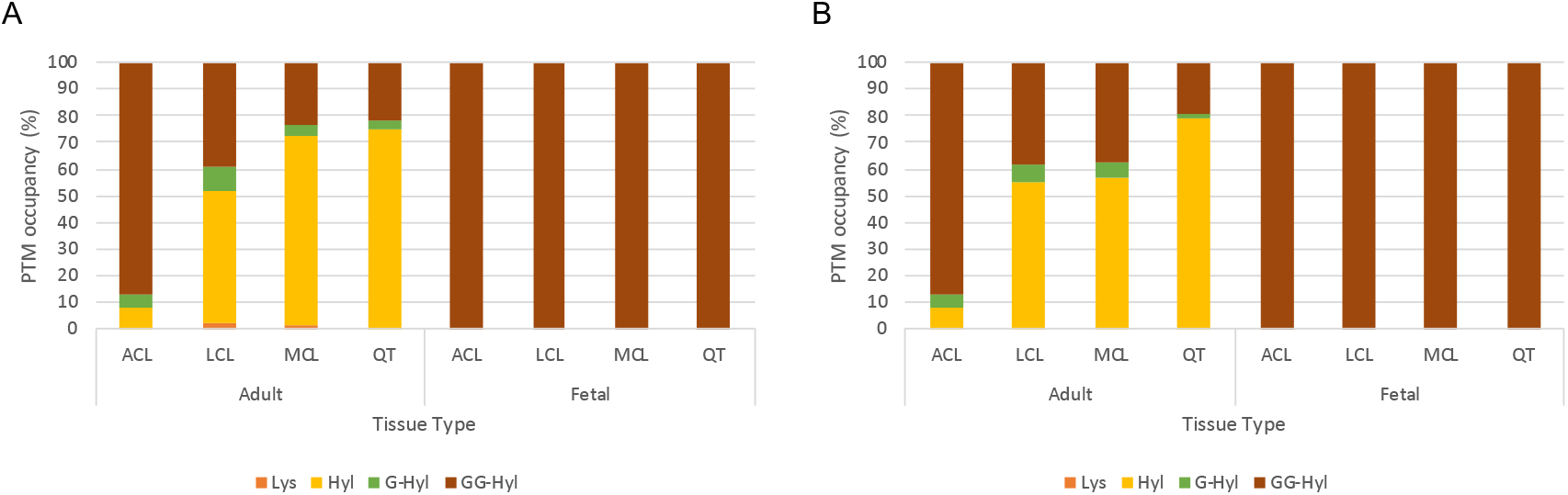
Post-translational variances in linear cross-linking α1(I)K87 and α2(I)K87 from type I collagens. Modifications on K87 from both A. α1(I) and B. α2(I) chains of type I collagen were measured in adult bovine tissues using mass spectrometry. Lysine modifications include unmodified lysine (Lys), hydroxylysine (Hyl), galactosyl-hydroxylysine (G-Hyl) and glucosylgalactosyl-hydroxylysine (GG-Hyl). The percentages were determined based on the ratios between the ion-current yields of each post-translational variant as previously described. See Table 2 for more details.

**Table 2.** Summary of post-translational variances in linear helical cross-linking lysines from type I collagens. Modifications on α1(I)K87, α2(I)K87, α1(I)K930 and α2(I)K933 from both α-chains of type I collagen were measured in adult and fetal bovine tissues using mass spectrometry. Lysine modifications include unmodified lysine (Lys), hydroxylysine (Hyl), galactosyl-hydroxylysine (G-Hyl) and glucosylgalactosyl-hydroxylysine (GG-Hyl). Mean percentages for K87 (n=3) and K930/933 (n=1-2) were determined based on the ratio the m/z peaks of each post-translational variant as previously described.

| **K87 hydroxylation (%)** |  |  |  |  |
| --- | --- | --- | --- | --- |
| Adult $\alpha 1(I)$ K87 | | | | |
|  | **Lys** | **Hyl** | **G-Hyl** | **GG-Hyl** |
| QT | 0 | 75±13 | 3±6 | 22±16 |
| ACL | 0 | 10±17 | 6±10 | 88±21 |
| LCL | 2.0±3 | 50±29 | 9±10 | 39.0±20 |
| MCL | 1±2 | 71±24 | 4±7 | 23±15 |
| Adult $\alpha 2(I)$ K87 | | | | |
|  | **Lys** | **Hyl** | **G-Hyl** | **GG-Hyl** |
| QT | 0 | 79±12 | 1±2 | 20±10 |
| ACL | 0 | 17±14 | 5±4 | 79±18 |
| LCL | 0 | 55±22 | 6±7 | 38±16 |
| MCL | 0 | 57±27 | 5±4 | 38±23 |
| Fetal $\alpha 1(I)$ K87 | | | | |
| QT | 0 | 0 | 0 | 100 |
| ACL | 0 | 0 | 0 | 100 |
| LCL | 0 | 0 | 0 | 100 |
| MCL | 0 | 0 | 0 | 100 |
| Fetal $\alpha 2(I)$ K87 | | | | |
| QT | 0 | 0 | 0 | 100 |
| ACL | 0 | 0 | 0 | 100 |
| LCL | 0 | 0 | 0 | 100 |
| MCL | 0 | 0 | 0 | 100 |

| **K930/K933 hydroxylation (%)** |  |  |
| --- | --- | --- |
| Adult $\alpha 1(I)$ K930 | | |
|  | **Lys** | **Hyl** |
| QT | 0 | 100 |
| ACL | 0 | 100 |
| LCL | 0 | 100 |
| MCL | 0 | 100 |
| Adult $\alpha 2(I)$ K933 | | |
|  | **Lys** | **Hyl** |
| QT | 0 | 100 |
| ACL | 0 | 100 |
| LCL | 0 | 100 |
| MCL | 0 | 100 |
| Fetal $\alpha 1(I)$ K930 | | |
| QT | 0 | 100 |
| ACL | 0 | 100 |
| LCL | 0 | 100 |
| MCL | 0 | 100 |
| Fetal $\alpha 2(I)$ K933 | | |
| QT | 0 | 100 |
| ACL | 0 | 100 |
| LCL | 0 | 100 |
| MCL | 0 | 100 |

The C-telopeptide cross-linking α1(I)K1030 residue was found to be partially hydroxylated in both fetal (65% hydroxylation) and adult (60% hydroxylation) tendons (Figure 5). In contrast, the C-telopeptide cross-linking lysine was almost fully hydroxylated in adult ligaments (~90%), while fetal ligaments, like tendon, exhibited only partial hydroxylation (~60%) (Figures 2 and 5). Ligaments from older fetuses were observed to have higher levels of C-telopeptide lysine hydroxylation than younger fetuses. This variability between fetal tissue can be appreciated by a high standard deviation in the calculated mean of percent hydroxylation (Figure 5). Similar to K930/K933, the hydroxylation state of the N-telopeptide cross-linking lysine is only detectable using bacterial collagenase digests (Figure 6). Fetal tissues were found to contain approximately 30-40% hydroxylysine at the N-telopeptide cross-linking site. Even in adulthood, this site remained about 50% hydroxylated in tendon (QT) (Figure 6C). However, the N-telopeptide cross-linking lysine was about 85% hydroxylated in adult ligaments (Figure 6A).

**Figure 4.**
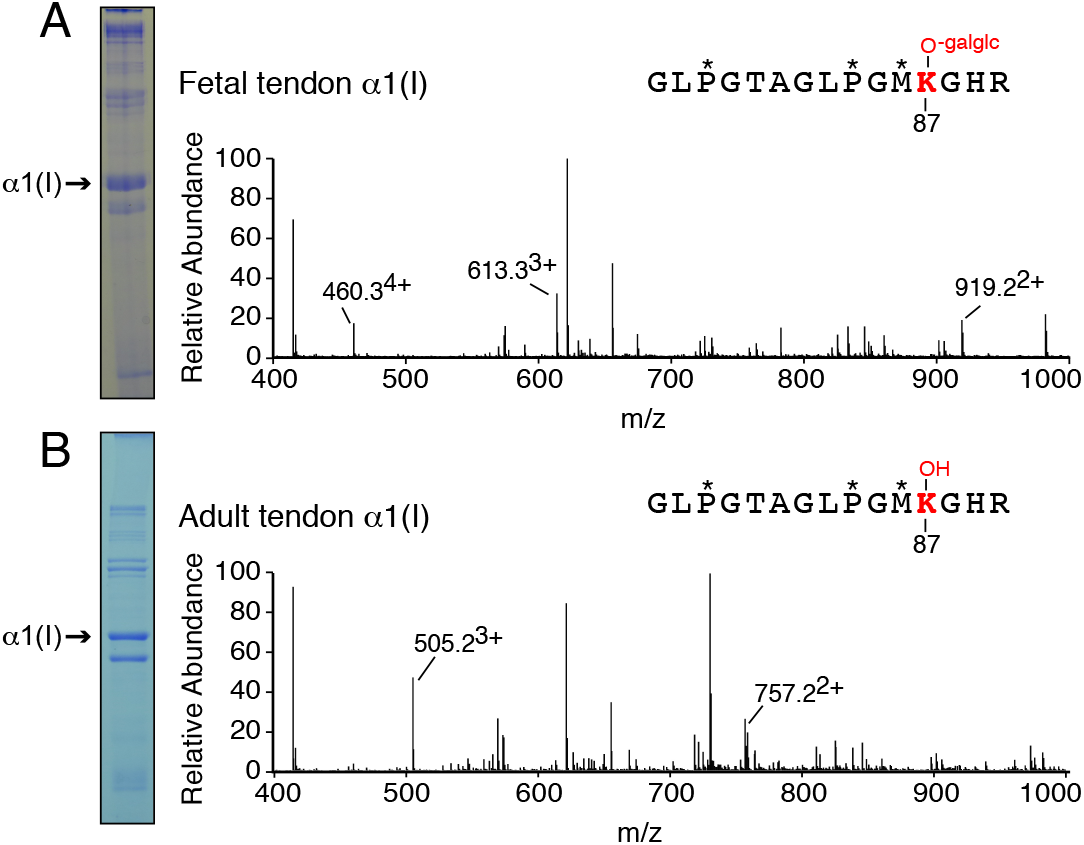
Decrease in glycosylation with age at type I collagen cross-linking residue (α1(I)K87) in quadriceps tendon. LC-MS profiles of in-gel trypsin digests of the collagen α1 (I) chain from fetal and adult bovine tendon. (A) Fetal bovine tendon shows complete glycosylation of α1(I)K87 (613.3^3+^) (B) The α1 (I)K87 residue from adult bovine tendon is hydroxylated but not subsequently glycosylated (505.2^3+^). The trypsin-digested peptide is shown with P* indicating 4Hyp, M* indicating oxidized methionine, galglc indicating glucosyl-galactosyl and K-OH indicating Hyl.

**Figure 5.**
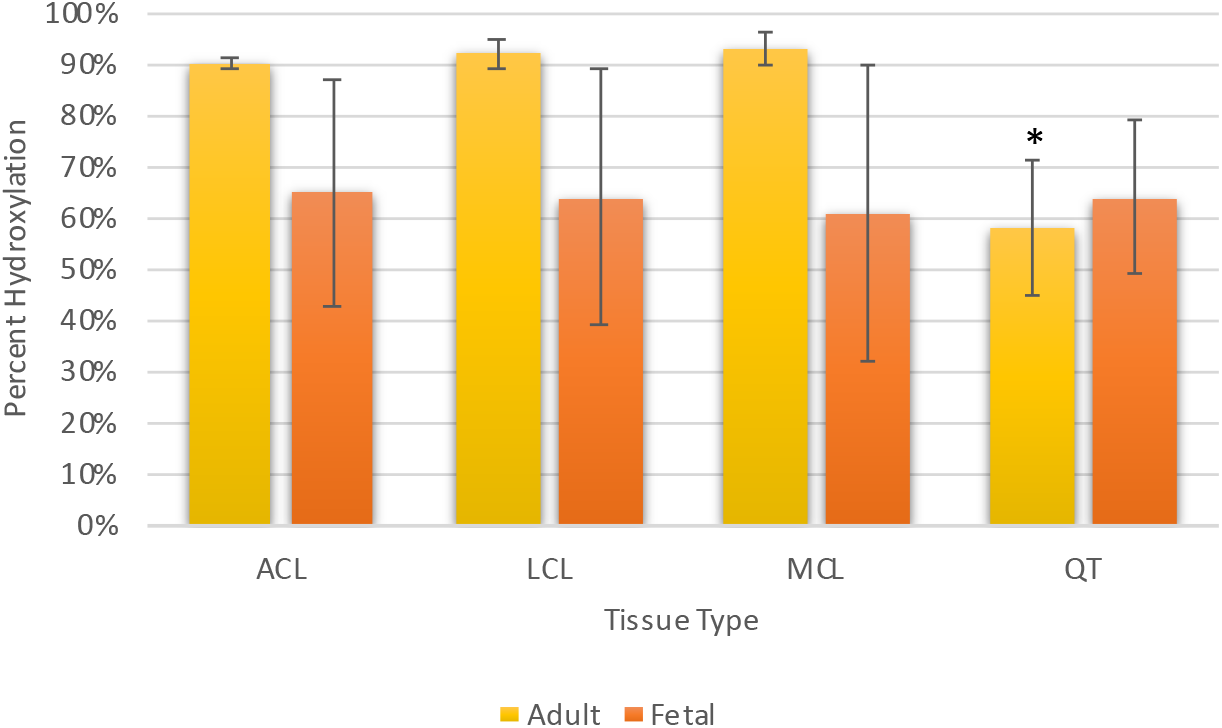
Increased hydroxylation of the linear C-telopeptide cross-linking lysine residue in adult ligaments. The C-telopeptide cross-linking lysine of type I collagen is a useful predictor of cross-linking quality across tissues. Percent hydroxylation is calculated from tryptic peptides using mass spectrometry (n=3). Note that lysine hydroxylation in adult QT is significantly less than adult ligaments; * p<0.05 by t-test assuming unequal variance.

**Figure 6.**
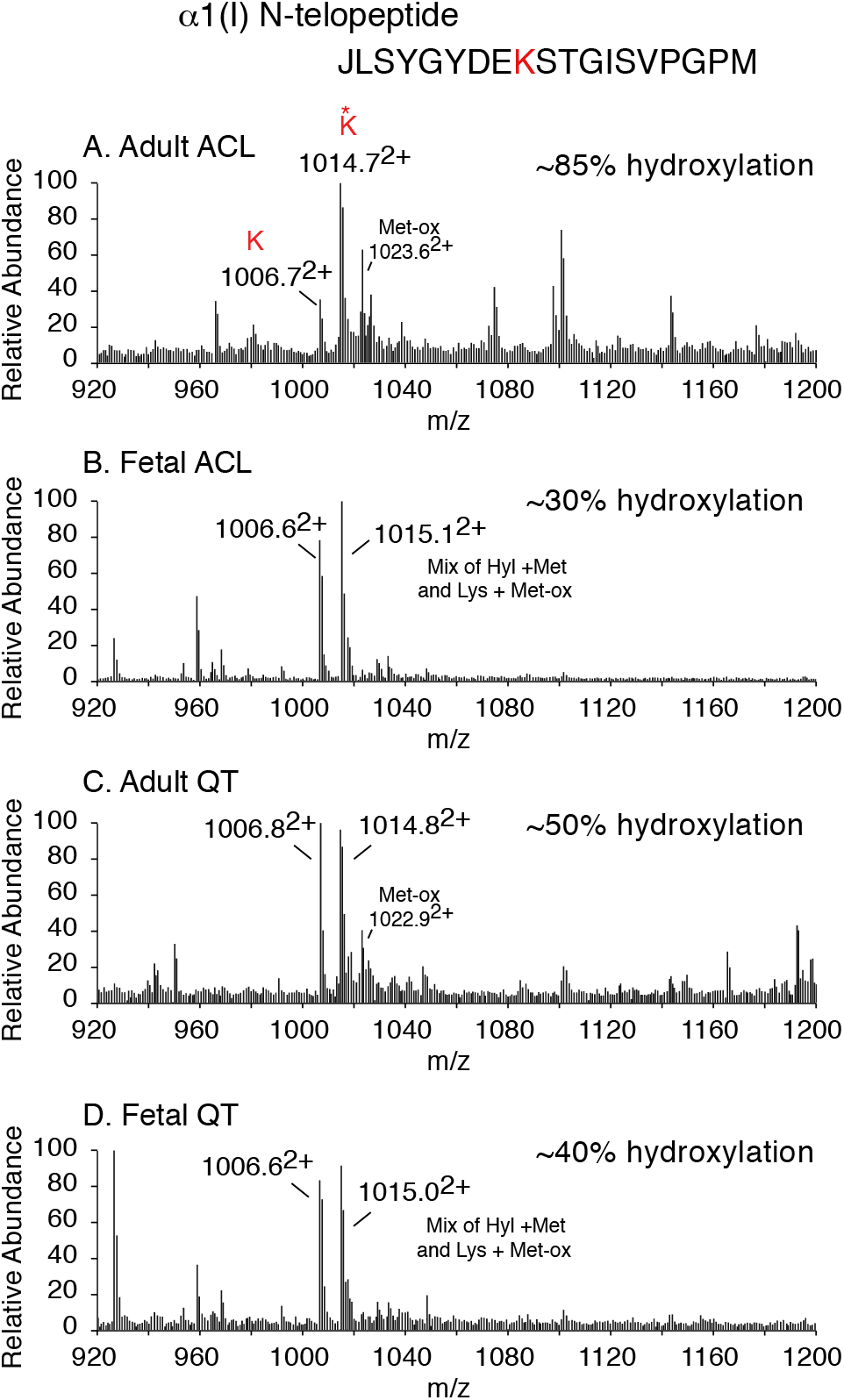
Ligament type I collagen has elevated N-telopeptide cross-linking lysine hydroxylation compared to tendon. LC-MS profiles of collagenase-digested adult ACL (A), Fetal ACL (B), Adult QT (C) and Fetal QT (D). (A) Adult ligaments had the highest level of hydroxylysine (~1015^2+^) at the N-telopeptide cross-linking site of all tissues tested. A small population of peptide containing hydroxylysine + oxidized methionine (~1023^2+^) was observed in adult tissues.

### Mature collagen cross-links are higher in ligaments than tendons

Despite more than half of the C-telopeptide cross-linking residues being hydroxylysine in fetal tendons and ligaments, pyridinoline cross-linking analysis revealed only negligible levels of HP in all fetal tissues tested (~0.06 mol/mol each). The relatively high levels of C-telopeptide hydroxylysine in fetal tissues accompanied with low levels of HP cross-links implies that fetal collagens contain mostly divalent cross-links that have yet to mature to HP. A partial C-telopeptide lysine hydroxylation profile is also consistent with fetal tendon and ligament collagens containing cross-links that use a different chemistry than the traditional hydroxylysine aldehyde cross-linking pathway, and/or containing less total cross-links than adult tissues [35,36]. The increase in telopeptide lysine hydroxylation in adult ligaments supports an increase in mature cross-links in mature ligament. Adult tendons, which were also found to have ~50% hydroxylation at α1(I)K1030, had ~10-fold higher levels of pyridinoline than fetal tissue (Figure 7). The ligamentous tissues had consistently higher levels of HP (~1.0 mol/mol) than tendon (0.5 mol/mol) in adult connective tissues, which is consistent with the higher levels of hydroxylation at α1(I)K1030 (Figures 5 and 6). Measurable levels of LP were not detected in any of the tissues tested. Another potential cross-linking product in type I collagen with partial hydroxylation of the telopeptide cross-linking lysine, as seen in the fetal tissues, is the trivalent cross-link pyrrole. The presence (or absence) of pyrroles in adult and fetal tendon and ligament was assessed using Ehrlich’s reagent (p-dimethylaminobenzaldehyde). Of all the tissues tested, only fetal tendon failed to turn pink after a 5 minute incubation in Ehrlich’s reagent, consistent with the absence of any pyrroles. Based on color intensity, fetal and adult ligament both contain pyrroles. Adult tendon produced the darkest pink color, potentially suggesting the highest level of pyrroles of the tissues tested (Figure 8) and consistent with the 50% telopeptide lysine hydroxylation (Figures 5 and 6). In this respect, tendon resembles bone [17].

**Figure 7.**
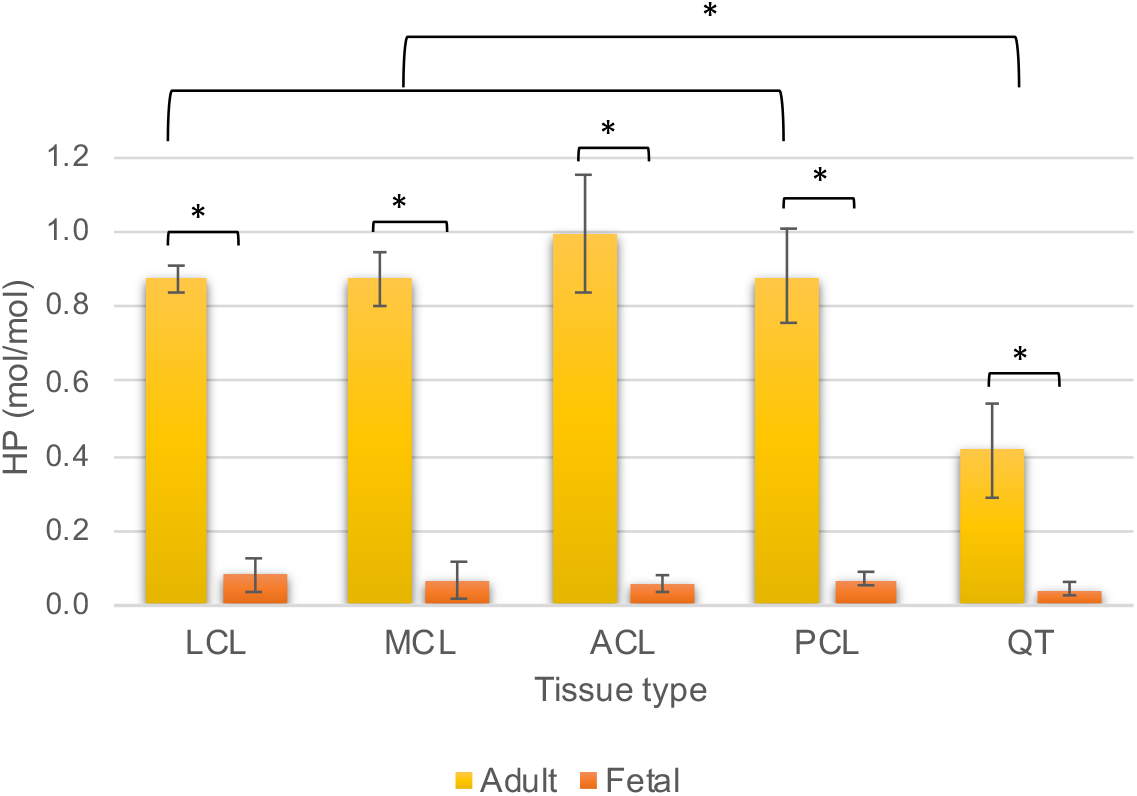
Increased pyridinoline cross-linking with increased tissue maturity. Concentration of hydroxylysine pyridinoline cross-linking residues in fetal and adult bovine tendon and ligaments expressed as moles/mole of collagen (n=3). (HP, hydroxylysylpyridinoline). None of the tissues contained measurable levels of LP (lysylpyridinoline). *p<0.05 by t-test assuming unequal variance.

**Figure 8.**
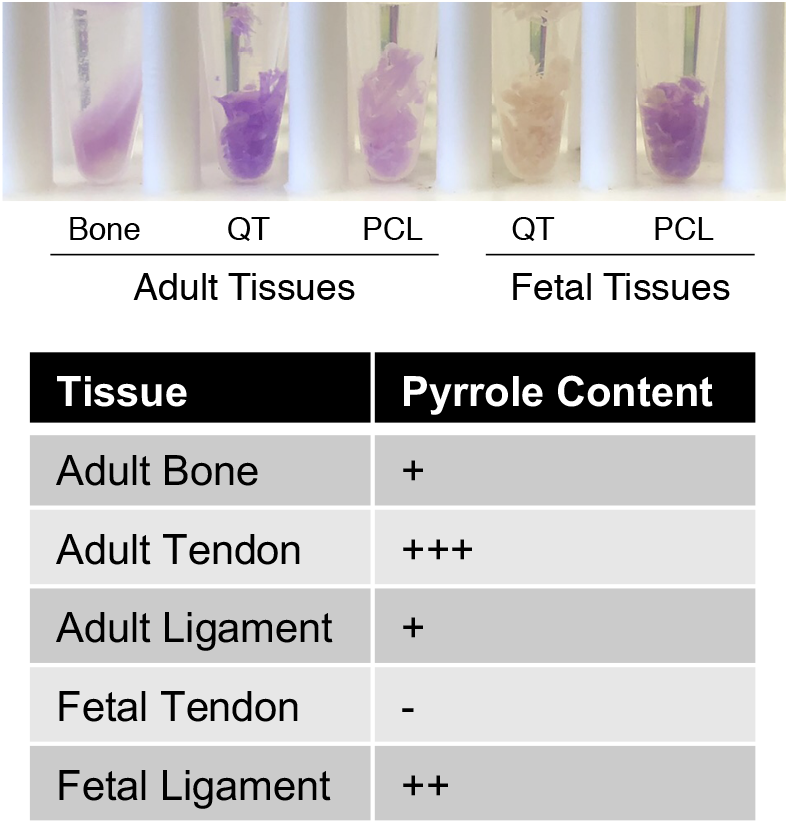
Tissue pyrrole content determined from Ehrlich reagent color change. Bovine tissues were minced and incubated in Ehrlich’s reagent for 5 minutes. No color change was detected in fetal tendon (n/d). The color reactions (n=2) were measured using online color detection software.

### Immature collagen cross-links are elevated in fetal tissues

Total collagen cross-links were quantified in acid hydrolysates of borohydride-reduced whole tissues. Cross-linking amino acid products were separated from free amino acids by organic solvent partition on a hydrated cellulose column and resolved by normal phase HPLC [37]. Total collagen cross-links are presented as ratios of ion-current yield relative to the ion current of HP from the same LCMS scroll (Table 3, Figure 9). The absolute mol/mol value of HP was calibrated from its independent HPLC fluorescence quantitation (Figure 7). Although this gives relative, not absolute, amounts of the other cross-links, the ratios give useful comparisons between tissues. It is evident from the cross-link ratios that histidinohydroxymerodesmosine (HHMD), a tetravalent cross-link formed by borohydride reduction between an N-telopeptide aldol, α1 (I)K930 and α1 (I)H932, is a prevalent cross-link in both the fetal tissues as well as adult tendon. As previously thought, both fetal tendons and ligaments are comprised of higher levels of immature cross-links hydroxylysinonorleucine (HLNL) and dehydrohydroxylysinonorleucine (DHLNL) than mature cross-links (HP) [38]. Important differences were also observed between adult tendon and ligaments. For example, the predominant cross-link measured in adult ligament was HP; however, adult tendon had significantly less HP yet appears to have more HHMD and pyrrole cross-links than adult ligament. Unfortunately, pyrrole cross-linking content cannot be measured using acid hydrolysis as the cross-link is acid labile.

**Figure 9.**
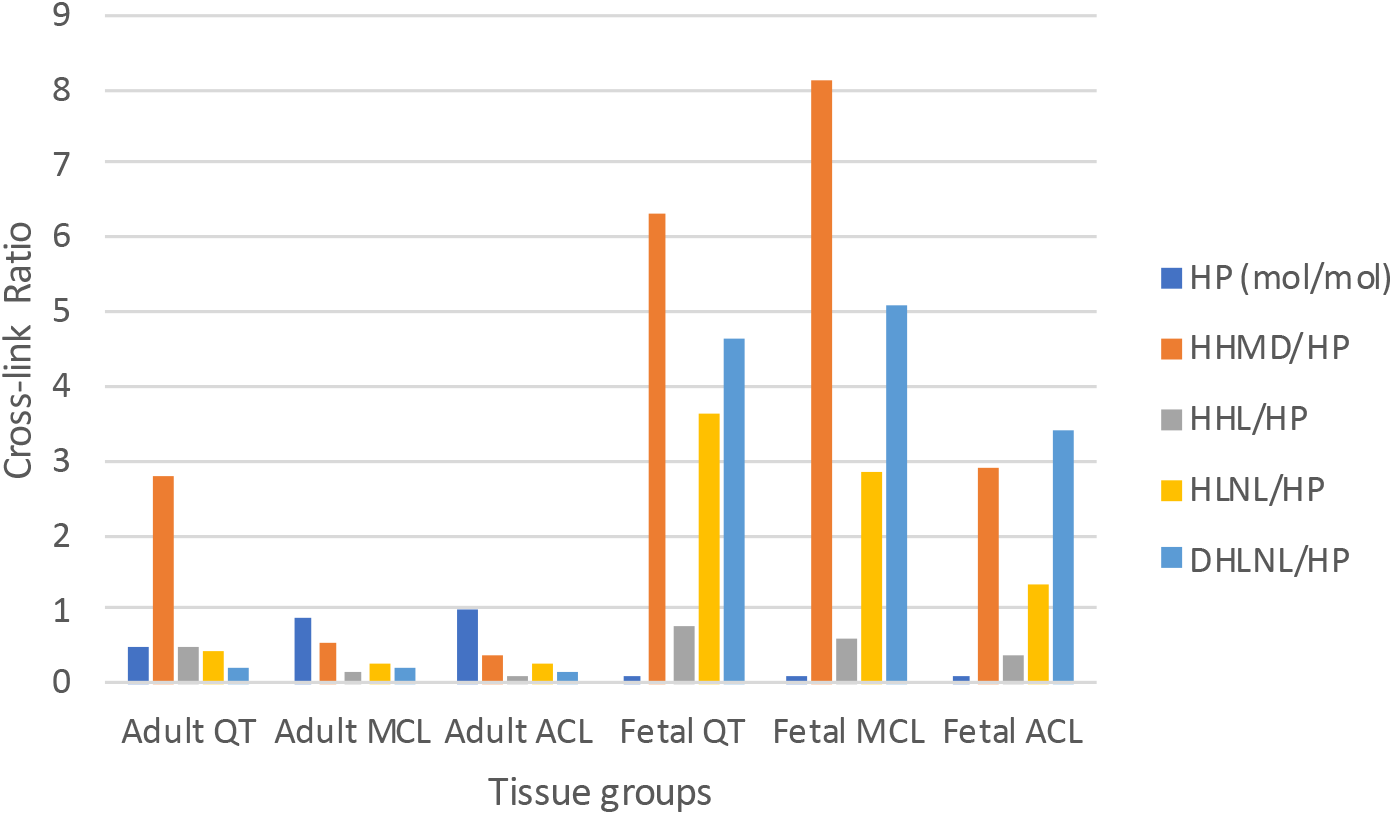
Comparison of bovine ligament and tendon total collagen cross-links. Borohydride reduced total collagen cross-links were determined from LC-MS profiles of tissue acid hydrolysates. Enriched cross-linking amino acids were resolved and analyzed by electrospray LC-MS on the LTQ XL using a Cogent 4 diamond hydride column. Cross-link populations are presented as ratio of HP (HP was quantitated using fluorescence) Values are the average of two analyses (n=2).

**Table 3.**
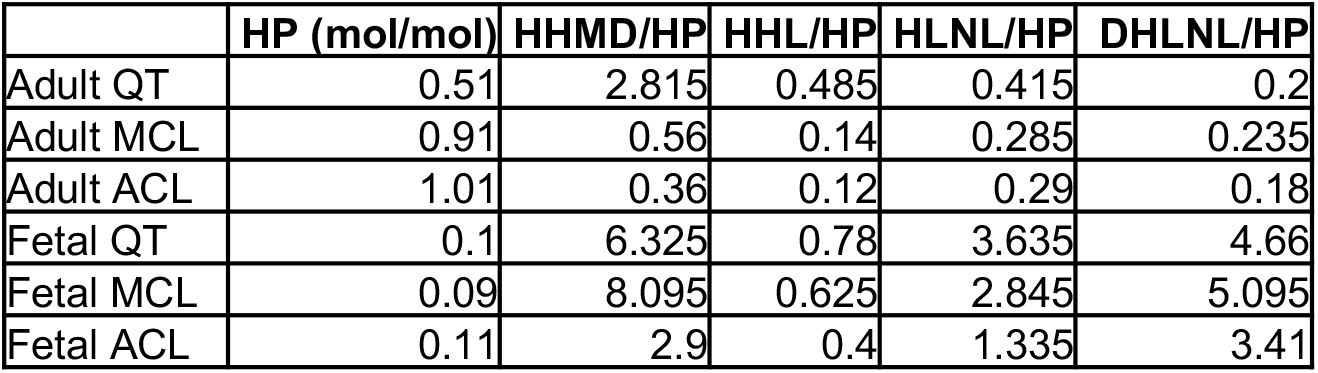
Total collagen cross-links relative to HP (values based on the intensity of each cross-link (m/z) in the total ion current; n=2). HP presented as mol/mol using HP calibrated by fluorescence HPLC.

|  | HP (mol/mol) | HHMD/HP | HHL/HP | HLNL/HP | DHLNL/HP |
| --- | --- | --- | --- | --- | --- |
| Adult QT | 0.51 | 2.815 | 0.485 | 0.415 | 0.2 |
| Adult MCL | 0.91 | 0.56 | 0.14 | 0.285 | 0.235 |
| Adult ACL | 1.01 | 0.36 | 0.12 | 0.29 | 0.18 |
| Fetal QT | 0.1 | 6.325 | 0.78 | 3.635 | 4.66 |
| Fetal MCL | 0.09 | 8.095 | 0.625 | 2.845 | 5.095 |
| Fetal ACL | 0.11 | 2.9 | 0.4 | 1.335 | 3.41 |

### Ultrastructural differences between tendon and ligament

Tendon (QT) and ligament (ACL) collagen fibrils were investigated by transmission electron microscopy (TEM) to assess potential ultrastructural differences associated with the unique cross-linking profile of each tissue. Thin sections of tendon and ligament were cut perpendicular to the long axis of the tissue in order to yield transverse collagen fibril transections. Overall, collagen fibril structure from tendon and ligament were quite similar as determined by TEM; however, several subtle distinctions were identified between the tissues (Figure 10A). For example, it was immediately apparent that ACL had more interfibrillar spacing than QT. This is consistent with ACL having less fibril density (number of fibrils/μm^2^) and fibril area fraction (percentage of ECM that is comprised of collagen fibrils) than QT (Figure 10C). A bimodal distribution of collagen fibril diameters was found in both tendons and ligaments, which were defined as small (30-90 nm) and large (140-200 nm) fibril populations (Figure 10B). A similar pattern in the bimodal distribution of fibril diameters was first observed in horse tendons and ligaments [39]. Tendon has a higher frequency of the small fibril population compared to ligament, whereas ligament has a higher frequency of the larger population compared to tendon (Figure 10B).

**Figure 10.**
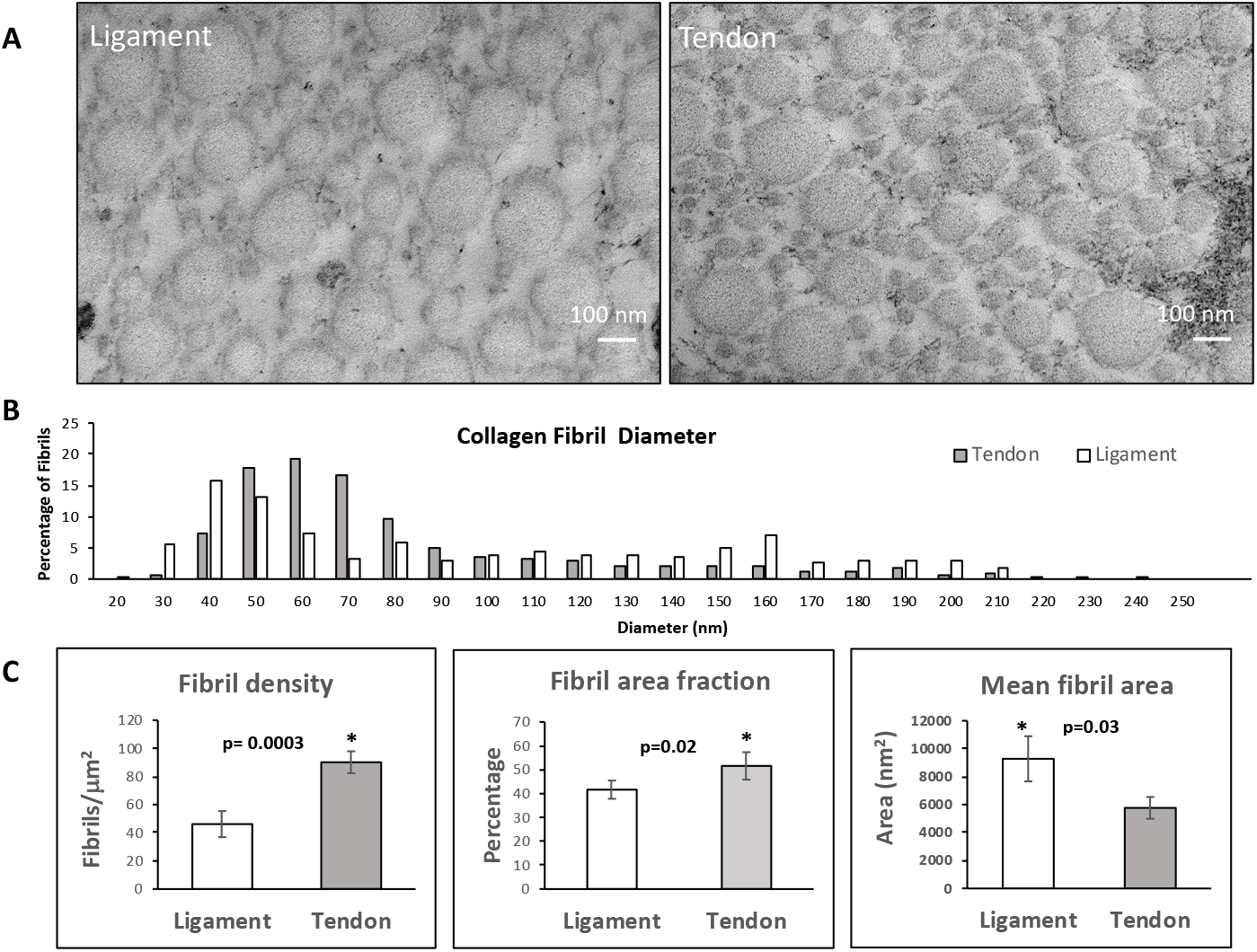
Transmission electron microscopy of collagen fibrils from bovine ligament and tendon. Representative transmission electron microscopy images showing slight distinctions between collagen fibrils between ligament (ACL) and tendon (QT) (A). Scale bar is 100 nm. Collagen fibril diameter plot illustrates bimodal distribution in both tissues (B). Fibril density, mean fibril area and fibril area fraction graphs illustrate several subtle distinctions between the tissues (C).

## Discussion

Tendons and ligaments both largely consist of type I collagen, but each has distinct evolutionary origins and functions. Tendons evolved early, before bone, from myosepta, sheet-like structures that transmitted primitive muscle force [40]. Although ligaments may have evolved from a similar origin, the primary function of most tendons is to transmit force needed for joint movement, whereas the primary function of most ligaments is joint stabilization and proprioception [5]. In this study, we attempted to distinguish between tendon and ligament with a focus on type I collagen post-translational modifications. It is becoming increasingly evident that the post-translational quality of fibril-forming collagens is central to understanding the functional and structural properties of a specific tissue [41]. Defining a unique molecular collagen phenotype between fetal and adult tissues may be important in understanding how injured fetal tendons are capable of tissue repair through effectively restoring the native collagen fibrillar architecture. Furthermore, evaluation of the distinct features of ligament type I collagen may provide insight into the post-translational adaptions that have evolved to enable the matrix to function in a non-vascularized environment.

Differences in post-translational modifications between tissues were first observed in the linear cross-linking K87 residues. For example, adult ACL contained significantly more K87 glycosylations than any other adult ligament collagen. In adult ACL, K87 from both α-chains was fully hydroxylated and predominantly glycosylated, specifically as glucosylgalactosyl-hydroxylysine. Of all the adult tissues tested, QT collagen had the lowest level of glycosylation at these cross-linking residues. It is not clear if the differences in glycosylation between intra-articular tissue (ACL, PCL) and more vascularized tissues (MCL, LCL, QT) imparts an advantage to these unique specialized tissues. Only negligible levels of non-modified lysine at K87 were detected in any adult tissue. In contrast, K87 from both α-chains was 100% glycosylated with glucosylgalactosyl-hydroxylysine in all fetal tissues tested. Although a specific role for hydroxylysine glycosylation in collagen cross-linking has yet to be fully elucidated, it has been proposed that the post-translational state of the helical-domain cross-linking lysines may serve as a regulator for normal collagen divalent aldimine cross-link chemistry [14]. In a mouse model, it was shown that reduction of hydroxylation and glycosylation at K87 resulted in a decrease in the level of intermolecular aldimine cross-links with C-telopeptide lysine aldehyde. Consequently, intramolecular aldol cross-links were increased relative to intermolecular aldol cross-links, which resulted in compromised material properties of the affected tissues [14,42]. Potential differences in collagen cross-linking that accompany tissue development were initially observed in the reduction of collagen α-chain extractability in adult tissues compared to fetal tissues as visualized by SDS-PAGE. This decrease in extractability supports the maturation of collagen cross-links from acid labile aldimines in fetal tissues to stable cross-links, such as pyridinolines, in adult tissues. An increase in pyridinoline content in type I collagen during tendon and ligament growth is consistent with collagen cross-linking changes that occur in cartilage [36] and bone development [43].

The high levels of hydroxylation (~60%) at the linear C-telopeptide cross-linking lysine across fetal tissues would support a higher level of pyridinoline and/or pyrrole than were found experimentally. Indeed, HP and pyrrole cross-links were notably absent in fetal bovine tendon, suggesting that the partially hydroxylated telopeptide lysines had yet to form these mature intermolecular cross-links in fetal collagens. It is tempting to speculate that the bulky sugar moiety at K87 might sterically hinder the formation of the mature cross-links. Borohydride-reduced fetal tendon was found to contain more HLNL, DHLNL and HHMD than fetal ligament. These findings are consistent with early amino acid analysis studies, which found fetal tendon to contain elevated levels of ketoimines, particularly DHLNL [38]. Unlike fetal tendon, fetal ligament also appears to contain some pyrrole cross-links. It should be noted that HLNL and borohydride-reduced lysinoketonorleucine are indistinguishable by mass (292 Da) using LC-MS. However, the high levels of hydroxylation of helical cross-linking lysines in fetal tissues combined with partial telopeptide lysine hydroxylation supports a prevalence of HLNL over the latter cross-link.

Adult tendon and ligaments could also be distinguished by the degree of telopeptide cross-linking lysine hydroxylation. The N- and C-telopeptide cross-linking lysine residues were over 80% hydroxylated in all adult ligament tissues, whereas adult and fetal tendon were only partially hydroxylated (~50%). This is consistent with adult ligaments having more mature HP cross-links than adult tendon. Partial hydroxylation at the telopeptide cross-linking lysines in adult tendon would also support tendon having higher levels of pyrrole cross-links, which is consistent with the Ehrlich’s reagent assay. In this respect tendon is similar to bone in having roughly 50% hydroxylation of telopeptide lysines [36], which may reflect a similar evolutionary origin. No significant differences were found between intra-articular ligaments and vascularized ligaments in regards to their C-telopeptide cross-linking lysine hydroxylation or HP content. Total cross-linking content from adult ligaments revealed that HP is the predominant cross-link in both intra-articular and vascularized tissues. By this analysis, adult tendon was unique in its cross-link composition in having almost three times more HHMD than any other cross-link. This is indicative of tendon using both the hydroxylysine aldehyde cross-linking pathway, which typically yields HP and/or LP, and the lysine aldehyde cross-linking pathway, which is prevalent in histidinohydroxylysinonorleucine (HHL) and/or HHMD cross-links. HHMD is an artifact of borohydride reduction, which, *in situ*, is a relatively labile cross-link comprised of an N-telopeptide aldol in Schff base linkage to Hyl [37]. Similarly, the Ehrlich reactive pyrrole from adult tendon is probably from N-telopeptide aldols linked to α2(I)K933, based on bone collagen chemistry [36,41,44].

Intermolecular collagen cross-linking is the definitive post-translational modification that imparts tensile strength on connective tissues. The unique cross-link profile of tendon (e.g. pyridinolines, pyrroles and aldols) may have evolved to support a long-lived tissue that must maintain tissue biomechanical properties during growth and remodelling phases. For example, the increased levels of labile intermolecular cross-links) may allow newly synthesized collagen molecules to fuse laterally to the growing fibril through labile cross-link breakages and reformation without compromising mechanical function [37]. A possible illustration of this cross-linking phenomenon is seen in the TEM images by the high frequency of small diameter fibrils in QT compared to ACL. Lateral fusion of these small fibrils to larger fibrils during tissue growth and remodelling would be facilitated by collagen molecules that accrue through less permanent cross-links. In other words, labile collagen cross-linking should enable the continual state of equilibrium needed on the surface of collagen fibrils to support tissue remodelling and growth, even in mature tendons. It is also likely that the distinct collagen cross-linking profile of tendon (and possibly ligament) is important for specialized tendon functions, such as interfibrillar tendon sliding to allow length growth with skeletal growth [45,46].

Type I collagen from tendon and ligaments was also surveyed for prolyl 3-hydroxylation content, as specific substrate sites for this rare modification have been uniquely localized to tendons [13,31]. Type I collagen from both tendon and ligaments shared a similar 3-hydroxyproline profile attributable to P3H1 [47] and P3H2 [33,48] activity. Another common feature between tissues was that fetal tissues have only minimal levels of prolyl 3-hydroxylation in type I collagen sites compared to adult tissues. The increase in 3-hydroxyproline previously associated with developing tendon [34], was also observed in across all bovine knee joint ligament collagens. It is clear from this work that fetal tendon and ligament cells have active P3H1, lysyl hydroxylase-1 (LH1), lysyl hydroxylase-2 (LH2) and glycosyltransferase enzyme networks. Fetal tissue has full lysyl 5-hydroxylation and glycosylation (glucosylgalactosyl-hydroxylysine) at cross-linking K87 residues from both α-chains. P3H2 and glycosyltransferase also appear to be uniquely regulated in ligament and tendon. Adult tendon cells, in particular, seem to experience a distinct decrease in glycosyltransferase activity, in addition to the accompanying increase in P3H2 activity. The result is adult tissues that have different collagen cross-linking profiles and likely unique structural and functional properties.

Why do only some ligaments heal? How do fetal tendons regenerate? It appears likely that the distinctive post-translational phenotypes of ligament and tendon collagens and how they change with development are related to these tissue-specific properties. It is becoming increasingly clear that as connective tissues develop and mature, their collagen molecular phenotype also changes. The unique molecular features observed between the collagens of these tissues likely highlights subtle yet distinct structural properties of each tissue. A greater understanding of collagen cross-linking during tendon and ligament development may provide insight into the pathobiology that leads to rupture and incomplete healing of mature tendon. Defining developmental variances in collagen post-translational modifications may be used diagnostically in the understanding of healthy tendon and ligament development and regeneration.

## Materials and methods

### Collagen extraction

Ligaments (ACL, PCL, MCL, LCL) and tendon (QT) were isolated from 18-month steer (adult) and fetal bovine knees purchased from Sierra for Medical Science (Whittier, CA). Intact type I collagen was solubilized from the tissues by acid extraction in 3% acetic acid for 24 hours at 4°C. Collagen α-chains were resolved by SDS-PAGE and stained with Coomassie Blue R-250. Total collagenase digests were resolved into peptide fractions on a C8 column by reverse-phase HPLC and monitored at 220nm.

### Pyridinoline cross-linking analysis

The pyridinoline cross-link content of collagen preparations was determined by HPLC after acid hydrolysis in 6 N HCl for 24 hrs at 108°C. Dried samples were dissolved in 1% (v/v) n-heptafluorobutyric acid and their reverse-phase HPLC lysyl pyridinoline (LP) and hydroxylysyl pyridinoline (HP) contents quantified by fluorescence monitoring as previously described [43].

### Pyrrole cross-linking analysis

The pyrrole cross-link content of collagen preparations was determined based on established protocols [17]. Briefly, equal amounts of tissues were finely minced (15 mg each) and resuspended in 6:1 (v/v)) Mili-Q H_2_O: 5% (w/v) p-dimethylaminobenzaldehyde in 4 M HClO_4_. The color reactions (n=2) were recorded after 5 minutes and measured using online color detection software (ColorMeter Free – color picker, vistech.projects).

### LC-MS peptide analysis

Lysine hydroxylation, glycosylation (glucosylgalactosyl-hydroxylysine and galactosylhydroxylysine), and prolyl 3-hydroxylation were quantified at specific sites in collagen α-chains as previously described [5,18]. Collagen α-chains were cut from SDS-PAGE gels and subjected to in-gel trypsin digestion. Samples were also digested with bacterial collagenase, with and without sodium borohydride reduction, and resolved by C8 reverse-phase HPLC prior to analysis by MS. Electrospray mass spectrometry was carried out on the trypsin and collagenase-digested peptides using an LTQ XL linear quadrapole ion-trap mass spectrometer equipped with in-line Accela 1250 liquid chromatography and automated sample injection [12]. Thermo Xcalibur software and Proteome Discoverer software (ThermoFisher Scientific) were used for peptide identification. Tryptic peptides were also identified manually by calculating the possible MS/MS ions and matching these to the actual MS/MS spectrum. Glycosylation and hydroxylation content in collagen α-chains differences were determined manually by averaging the full scan MS over several minutes to include all the post-translational variations of a given peptide. Protein sequences used for MS analysis were obtained from the Ensembl genome database.

### LC-MS cross-link analysis

Freeze-dried bovine tendon and ligament samples were reduced with sodium borohydride and hydrolyzed in 6 N HCl at 108°C for 24 h. Cross-linking amino acids were enriched from hydrolysates by partition on a hydrated cellulose column as previously described [37,49]. Briefly, the hydrolyzed sample was loaded on a cellulose (CF1, Whatman) column in butanol/ water/acetic acid (4:1:1, v/v/v). Free amino acids were removed by washing the column with butanol/water/acetic acid solution. Cross-linking amino acids were eluted with water and freeze-dried for mass spectral analysis. Electrospray LC-MS/MS was performed on the LTQ XL using a Cogent 4 diamond hydride column (15 cm x 1 mm; Microsolv Technology, 70000-15P-1) eluted at 50 μl min. The LC mobile phase consisted of buffer A (0.1% formic acid in MilliQ water) and buffer B (0.1% formic acid in 80% acetonitrile) [37].

### Transmission electron microscopy (TEM)

Bovine anterior cruciate ligament and quadriceps tendon samples were dissected from the knee joint and prepared for TEM using a modification of the procedure we have published before [50]. Mid-section samples from both tissues were rinsed 3X in PBS pH 7.4 at 4°C for 2 h. Samples were fixed in 2.5% paraformaldehyde and 0.25% glutaraldehyde in 0.2 M sodium phosphate buffer pH 7.4 for 24 hours. Post-fixation was carried out using 2% osmium tetraoxide in 0.2 M sodium phosphate buffer, pH 7.4. The samples were then dehydrated and embedded in plastic and cured overnight. Ultrathin transverse sections (70–90 nm) were cut, placed on 150-mesh copper grids, and stained with 4% uranyl acetate and Reynold’s lead citrate. All sections were examined and photographed on a FEI Tecnai Spirit Bio-Twin transmission electron microscope, fitted with a Hamamatsu ORCS HR camera. For each sample a minimum of 4 different regions were viewed. An unbiased sampling of each region was performed. A magnification of 30000X was used for collagen fibril area, collagen fibril area fraction and collagen fibrils per μm^2^. All measurements were made using ImageJ software (NIH freeware, http://rsb.info.nih.gov/nih-image) [51]. Areas of collagen fibrils in 7.28 μm^2^ of ligament and tendon was measured. Mean fibril area was calculated from average collagen area and number of fibrils counted. Collagen area fraction was calculated using total collagen area measured in the total area sample. Fibril diameter was calculated from fibril area assuming circularity (1/4 × π × diameter^2^). Data are presented as mean ± SD. For statistics, the two-sample assuming unequal variances t-Test was used and alpha was set to 0.05. p values less than 0.05 were considered to be significant.

## Abbreviations

(ACL): Anterior cruciate ligament
(PCL): posterior cruciate ligament
(LCL): lateral collateral ligament
(MCL): medial collateral ligament
(QT): quadriceps tendon
(HP): hydroxylysine pyridinoline
(LP): lysine pyridinoline
(HLNL): hydroxylysinonorleucine
(DHLNL): dehydrohydroxylysinonorleucine
(HHL): histidinohydroxylysinonorleucine
(HHMD): histidinohydroxymerodesmosine
(P3H1): prolyl 3-hydroxylase 1
(P3H2): prolyl 3-hydroxylase 2
(LC): liquid chromatography
(MS): mass spectrometry

## Author contributions

Conceptualization, D.M.H., D.R.E.; Supervision, D.M.H., D.R.E.; Data Collection and Analysis, D.M.H., M.A., M.A.W. R.J.F.; Formal analysis; D.M.H., R.J.F., D.R.E.; Roles/Writing - original draft, D.M.H.; Writing - review & editing D.M.H., R.J.F., D.R.E.; Funding acquisition, D.M.H., R.J.F., D.R.E. All authors intellectually contributed and provided approval for publication.

## Acknowledgements

This work was supported by the National Institutes of Health (NIH) Grants: (NIAMS) AR037318 and (NICHD) HD070394 to DRE; (NIAMS) AR057025 to RJF; (NIA) AG065605 to DMH. The authors would like to thank Jennifer K. Swicord, Electron Microscopy Supervisor, Department of Laboratory Medicine and Pathology, University of Washington Medical Center for expert technical assistance.

